# B lymphocytes in Treatment-Naïve Pediatric Lupus Patients are Epigenetically Distinct from Healthy Children

**DOI:** 10.1101/2022.09.23.509218

**Authors:** Joyce S Hui-Yuen, Kaiyu Jiang, Susan Malkiel, B Anne Eberhard, Heather Walters, Betty Diamond, James N. Jarvis

**Author notes:** **Correspondence**: Joyce S Hui-Yuen MD, MS, Associate Professor, Center for Autoimmune, Musculoskeletal, and Hematopoietic Diseases Research Feinstein Institute for Medical Research, 350 Community Drive, Manhasset NY 11030.

## Abstract

**Background:** Systemic lupus erythematosus (SLE) is a complex disease likely triggered by gene-environment interactions. We have shown that most of the SLE-associated haplotypes encompass genomic regions enriched for epigenetic marks associated with enhancer function in neutrophils, and T and B cells, suggesting that genetic risk is exerted through altered gene regulation. Data remain scarce on how epigenetic variance contributes to disease risk in pediatric SLE (pSLE). We aim to identify differences in epigenetically-regulated chromatin architecture in treatment-naïve pSLE patients compared to healthy children.

**Methods:** Using the assay for transposase-accessible chromatin with sequencing (ATACseq), we surveyed open chromatin in 8 treatment-naïve pSLE patients, with at least moderate disease severity, and 5 healthy children. We investigated whether regions of open chromatin unique to pSLE patients demonstrate enrichment for specific transcriptional regulators, using standard computational approaches to identify unique peaks and a false discovery rate of <0.05. Further analyses for differential transcription factor binding, histone modification enrichment, and variant calling were performed using multiple bioinformatics packages in R and Linux.

**Results:** There were 30,139 differentially accessible regions (DAR) identified unique to pSLE B cells, of which 64.3% are more accessible in pSLE than B cells from healthy children. Many of these DAR are found in distal, intergenic regions, and are enriched for enhancer histone marks (p=0.027). When we compared B cells from pSLE patients to those of untreated adults, we found more regions of inaccessible chromatin, and fewer DAR within 10-100kb of known SLE haplotypes. In pSLE B cells, 65.2% of the DAR are located within or near known SLE haplotypes. Further analysis revealed enrichment of several transcription factor binding motifs within these DAR that may regulate genes involved in the pro-inflammatory responses and cellular adhesion.

**Conclusions:** This is the first report describing differences in chromatin architecture between pSLE patients and healthy children. We demonstrate an epigenetically-distinct profile in pSLE B cells when compared to those from healthy children and adults with lupus, indicating that pSLE B cells are predisposed for disease onset and development. Increased chromatin accessibility in non-coding genomic regions controlling activation of inflammation and the immune response suggest that transcriptional dysregulation by regulatory elements that control B cell activation plays an important role in the pathogenesis of pSLE.

## Introduction

Systemic lupus erythematosus (SLE) is a complex disease thought to be triggered by gene-environment interactions. It is often marked by periods of disease flare and remission, and optimal treatment remains challenging. Pediatric-onset SLE (pSLE) accounts for up to 20% of all patients with SLE and can have a more severe and debilitating course than adult SLE; pediatric patients experience a shorter time to first intensive care admissions and are typically treated with multiple immunosuppressive medications to achieve disease control (1,2). We have shown that much of the SLE genetic risk identified in several genome-wide association studies (GWAS) lies in functional, non-coding (3) regions of the genome that are responsible for the regulation of transcription.

Functional elements that regulate and coordinate transcription are typically found in regions of ‘open’ chromatin (4,5) and are marked by specific epigenetic signatures consisting of specific histone marks (6,7). These histone marks are abundant in non-coding regions of the genome, affect other functional elements, and have promoter or repressor properties, leading to altered chromatin accessibility and ultimately affecting gene expression. Coit et al (8,9) have shown that gene expression and regulation are specific to distinct cell types, and that cell-specific expression is a feature of chronic diseases such as SLE. Dozmorov et al (10) demonstrated that adult SLE B cells express different genes than a population of monocytes from the same patients. It has been well-established that B cells are of central importance in disease pathogenesis in SLE, and contribute toward disease flares (reviewed in 11). These data also suggest that SLE may develop secondary to aberrant transcriptional and epigenetic control in B cells, resulting in the cells’ inability to coordinate and regulate transcription across the genome.

One way to investigate epigenetic changes in any chronic disease is to examine chromatin architecture, including accessible regions, using assays such as DNAse I hypersensitivity assays or assays for transposase-accessible chromatin with sequencing (ATACseq). Pioneered by Buenrostro et al (12), ATACseq allows for the surveying of the epigenome for differences in chromatin architecture and accessibility. Traditional methods such as DNAse I hypersensitivity assays typically require a large number of cells (often over 1 x 10^8^) to achieve adequate sequencing depth. Initially ATACseq required as few as 50,000 cells; the advent of newer sequencing technologies now allow ATACseq at the single cell level (single cell ATACseq, 13), making ATACseq a more feasible assay to generate high quality data for pSLE research.

To date, only one other group has used ATACseq to investigate epigenetic influences on disease development in adult SLE B cells: Scharer et al (14) described disordered chromatin architecture in adult SLE B cells when compared to healthy adult B cells. In particular, they noted increased chromatin accessibility for transcription factors important in B cell activation and differentiation in SLE B cells, in contrast to increased accessibility to transcription factors important in processes associated with regulation of transcription in healthy adult B cells. Gensterblum et al (15) used ATACseq to investigate chromatin accessibility in a newly defined T cell subset thought to be important in predicting disease activity in adult SLE. Data remain scarce on how epigenetic variance contributes to disease risk in pediatric SLE (pSLE) or how epigenetic features of the genome are influenced by genetic variants. We aim to identify differences in epigenetically-regulated chromatin architecture in treatment-naïve pSLE patients compared to healthy children.

## Methods

### 2.1 Patient enrollment

We obtained approval to conduct this prospective, observational study from the Northwell Health Institutional Review Board (IRB# HS16-0017) and informed consent from all enrolled patients and/or parents. Assent was obtained from children between the ages of 7 and 17 years in accordance with IRB policy. All patients seen at the Pediatric Rheumatology clinic at Cohen Children’s Medical Center between February 2016 and April 2018 with pediatric-onset lupus (diagnosed prior to their 19^th^ birthdays), who had not received prior treatment for lupus, were eligible for enrollment into this prospective, observational study. All enrolled patients fulfilled the 1997 ACR classification criteria (16), and had a disease activity index score (SLEDAI) of over 4 at diagnosis, indicating at least moderate disease severity (17). Healthy children were identified and enrolled from our General Pediatric and Adolescent Medicine clinics. Children were excluded if they had another autoimmune or chronic inflammatory disease, were obese (body mass index above 30), were acutely ill with fever or on antibiotics for a recent infection, or on corticosteroids for any reason.

### 2.2 Patient demographics

Ten newly diagnosed, treatment-naïve patients with pSLE and five healthy children were enrolled in this study. There were 7 female (70%) and 3 male (30%) patients in the pSLE cohort, ranging from 7-17 years of age. There were 4 healthy females (80%) and 1 healthy male (20%), ranging from 9-18 years of age. In the pSLE cohort, 50% were Black, 20% White, and 30% Asian, with 50% identifying as Hispanic ethnicity. Healthy children were 60% Black and 40% White, with 40% identifying as Hispanic ethnicity.

In the pSLE cohort, the mean age of disease onset was 12.5 (range 7-17) years of age, and mean SLEDAI 13 (range 6-24). All patients had a SLEDAI over 4, indicating at least moderate disease activity at time of enrollment. Lupus manifestations of rash and arthritis were present in 90% of the patients, lymphopenia in 70%, and neurologic disease in 20%. Biopsy-proven lupus nephritis was found in 50% of the patients. All patients had serologies positive for anti-nuclear autoantibodies (ANA) and anti-double stranded DNA (dsDNA) autoantibodies, which are specific for a diagnosis of lupus. All patients also had low complement levels, which are a marker for active disease.

### 2.3 B cell isolation and enrichment

At time of enrollment, 20ml of whole blood was collected from each patient (treatment-naïve lupus patients and healthy children) and processed within four hours of arrival in our laboratory. B cells were obtained by negative selection using the Stem Cell Technologies (Vancouver, BC, Canada) B cell enrichment kit (product #19054), using anti-CD3 (BioLegend #300469), anti-CD14 (BioLegend #301853), anti-CD16 (BioLegend #302059), and anti-CD19 (BioLegend #302212) antibodies. Cells obtained in this manner were 95% CD19+ when assessed by flow cytometry.

### 2.4 Assays for transposase-accessible chromatin with sequencing (ATACseq) preparation and statistical analysis

After B cell enrichment, 50,000 cells were transferred into sterile DNAse/RNAse free Eppendorf tubes on ice and washed three times with cold phosphate buffered saline (PBS). Resulting pellets were then kept at −80°C until ready for shipment to the Jarvis laboratory for ATACseq library preparation.

ATACseq consists of 3 major steps: preparation of nuclei, insertion of Tn5 transposases and DNA purification, and PCR amplification across sequencing adapters. We used the ATACseq protocol described by Buenrostro et al (12). Briefly, nuclei were prepared by spinning 50,000 CD19+ B cells at 500*g* for 5 minutes, then washed using 50ul of cold phosphate buffered saline (PBS) and spun again at 500*g* for 5 minutes. Cells were lysed using cold lysis buffer (10nM Tris-Cl, pH 7.4, 10mM NaCl, 3mM MgCl2 and 0.1% IPEGAL CA-630). Immediately post-lysis, nuclei were spun at 4°C at 500*g* for 10 minutes and re-suspended in the transposase reaction mix (25ul 2X TD buffer, 2.5ul transposase (Illumina) and 22.5ul nuclease-free water). The transposition reaction was carried out at 37°C for 30 minutes; immediately after, the Qiagen Minelute kit was used to purify the sample. Post-purification, library fragments were amplified using 1X NEBnext PCR mastermix and 1.25uM PCR primer1 and barcoded PCR primer2. The following thermocycler conditions were used: 5 minutes at 72°C, 30 seconds at 98°C, cycling for 10 seconds at 98°C, 30 seconds at 63°C, and 1 minute at 72°C. We monitored the PCR reaction using quantitative PCR to reduce GC and size bias in the reactions by amplifying the full libraries for 5 cycles, then adding 5ul of the PCR reaction to 10ul PCR cocktail with SyBR green at a final concentration of 0.6x. To determine the additional number of cycles needed for the remaining 45ul reaction, we ran the 15ul reaction for 20 cycles, and stopped amplification prior to saturation. We then plotted linear Rn versus cycle to determine the cycle number corresponding to 1/3 of the maximum fluorescent intensity. AMPure XP beads were used to purify the libraries, yielding a final library of 17.5ul. Libraries were then sent to the Genomics and Bioinformatics Core facility at the University at Buffalo for 50bp paired-end sequencing on the Illumina HiSeq 2500 platform.

### 2.5 Data analysis

Conventional computational approaches were used for peak-calling. In brief, raw data (FASTQ) files had adapters trimmed using Trim Galore (18), and quality control using FASTQC (19). Files were then aligned to the human genome (hg19) using BBMap, a splice-aware aligner for DNA and RNA sequencing reads, with MAPQ>30, and sorted using Samtools (20,21). Only uniquely-mapped and non-redundant reads were used for downstream analysis. Peak calling was performed using Model-Based Analysis for ChIP-seq (MACS2) using its default settings (22). Differential accessibility analysis was performed using DiffBind (23) and centered around peak summits, extending 1000 base pairs in each direction. Peaks with false discovery rate (FDR) <0.05 were considered significantly differentially accessible between pSLE and healthy children. Histone modification analysis was performed using the Roadmap/ENCODE database and gene ontology analysis for biological function with the GO database. Motif detection was performed using PScan-ChIP and HOMER (24,25). Peaks were annotated using ChIPpeakAnno and ChIPseeker. Nucleosome positioning analysis was performed using NucleoATAC in Python and NucHunter, a Java-based application. Gene ontology enrichment analysis was performed using the publicly-available Gene Ontology enRIchment analysis and visualization tool (GORILLA) (26), and further reduced using the Reduce Visualize Gene Ontology tool (REVIGO) based on semantic similarity measures (27).

### 2.6 Comparison of ATACseq data from adult and pediatric B cells

We compared ATACseq data from adult SLE and pSLE B cells. In brief, we downloaded and queried ATACseq data from CD19+ B cells of 15 adult lupus patients and 51 healthy adults from the publicly-available NCBI Sequence Read Archive (SRA, dataset GSE118256, 14). No further annotation of these patients was available. Quality inspection of the downloaded FASTQ files was performed using FASTQC (19). The reports for all FASTQ files were combined (by condition: SLE and healthy) using MULTIQC (28). No red flags requiring remediation were found. Alignment to the human GRCh38 reference genome was performed using Bowtie2 with its default parameters (29). Major SNPs from 1K-Genomes were included with the reference genome (30). Post-alignment, SAM files were converted to BAM files, sorted, and indexed using Samtools (21). The resulting BAM files were shifted using deepTools2 (left-most fragments were shifted 4 bases to the right, and right-most shifted 5 bases to the left) (31).

Similar methodology was used to call peaks and compare DARs between adult and pediatric SLE B cells as was used to identify DARs between pediatric patients and healthy children. DARs from the adult SLE patients were subtracted from pSLE DARs, using Bedtools, to focus on the differences attributable to disease pathology (32). Motif analysis was again performed using PScan-ChIP and HOMER (24,25) and annotated with ChIPseeker.

## Results

### 3.1 B cells from pediatric lupus patients exhibit a unique chromatin architecture

Differential peak analysis identified 30,139 uniquely accessible sites in pSLE patients (Figure 1) when compared with the B cells of healthy control children. Of these sites, 64.3% are more accessible in pSLE patients than healthy children (Figure 1c), but there were also regions that demonstrated decreased accessibility in the B cells from pSLE patients, when compared to those from healthy children.

**Figure 1.**
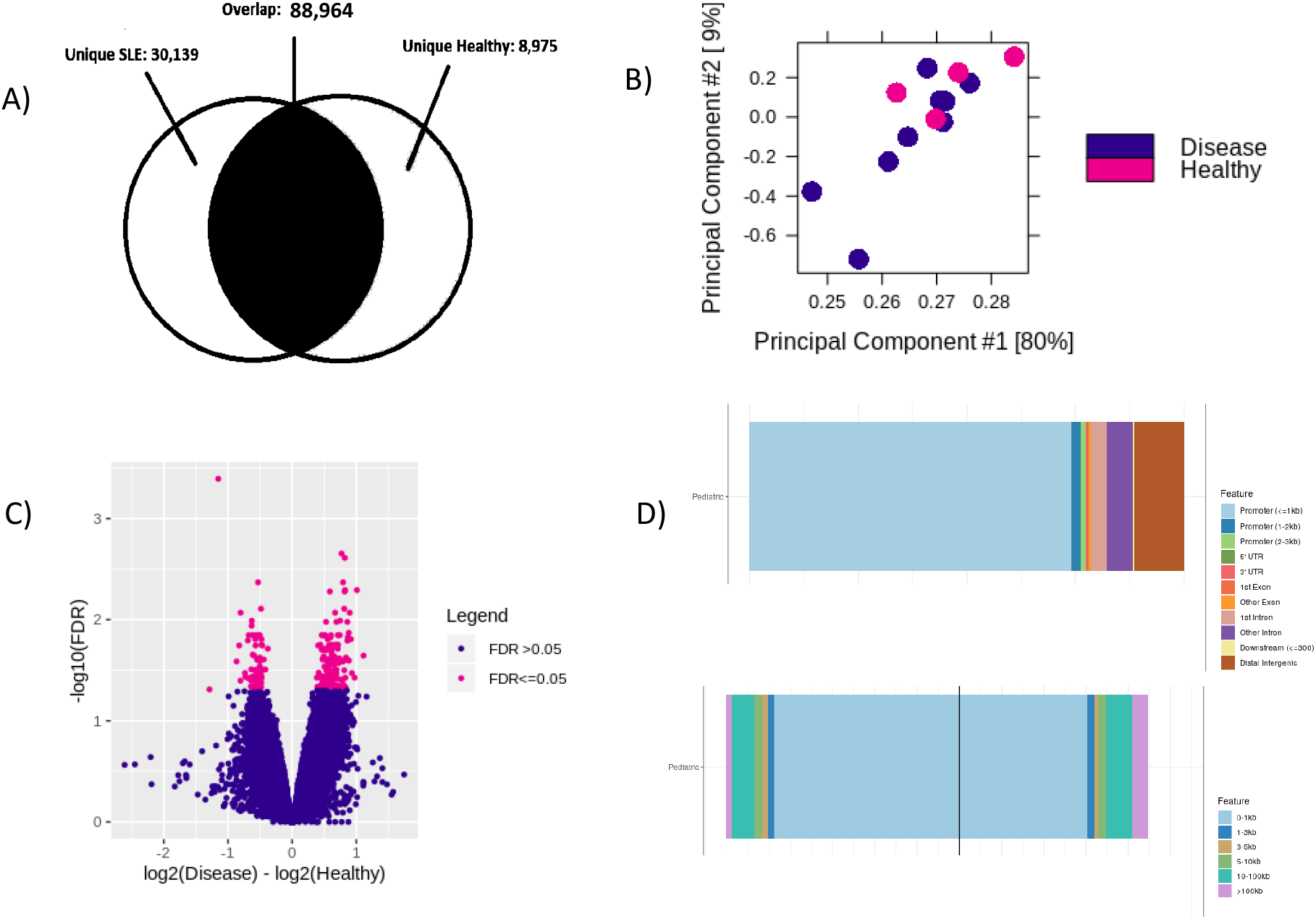
Unique chromatin architecture in pediatric-onset lupus B cells compared to B cells from healthy children. Peaks distinct from B cells from healthy children tend to be located distally from transcription start sites. a) Venn diagram showing unique peaks in pediatric lupus patients (30,139) and healthy children (8,975). b) Principal component analysis plot showing distribution of pediatric lupus (darker points) and healthy children (lighter points) samples. Note that the lupus samples in closer proximity to healthy children samples have milder disease severity. c) Differentially accessible regions in pediatric lupus B cells compared to healthy children (false discovery rate <0.05). Lighter plotted points to the right of 0 on the x-axis indicate increased chromatin accessibility in pediatric lupus patients. Lighter plotted points to the left of 0 on the x-axis indicate potentially less accessible chromatin in pediatric lupus patients. d) Top histogram showing that over 75% of unique peaks in pediatric lupus patients are located 10-100kb or more from transcription start sites. Bottom histogram showing that the peaks that are located around transcription start sites are enriched for transcription factor binding loci.

There were 50.34% of the differentially accessible regions (DAR) in pSLE found in distal, intergenic regions of the genome; only 7.7% of the DAR are located directly in promoters or exons. Further analyses of these open regions using ChipEnrich revealed that 65.4% of these unique peaks are located between 10-100kb and above from the nearest transcription start site (Figure 1d), implying that many transcription factors may be acting within distal enhancers to regulate transcription.

We previously identified 46 single nucleotide polymorphisms (SNPs) from GWAS in adults with SLE that lie in distal intergenic regions and may regulate transcription (3). We compiled a list of another 325 SNPs from more recent GWAS in adult SLE that are enriched for enhancer function and thus may regulate transcription (33–35). Using NucleoATAC to interrogate broad peaks obtained from pSLE patients, and Bedtools intersect to identify nucleosome positions that overlap with DAR, generated 158 nucleosome positions that lie within 10-100kb of SNPs used to tag SLE risk loci identified by GWAS and our previous study (3,33–35). We then used SNiPA (single nucleotide polymorphisms annotator) (36) to determine the haplotypes defined by these SNPs (r^2^ <0.9). There were 65.2% of the DAR in pSLE patients located within these known SLE haplotypes. Local or cis expression quantitative trait loci (eQTLs) were found in 43.2% of the SLE haplotypes containing DAR from pSLE patients; similarly, 34.3% of these haplotypes contained trans eQTLs.

When the DAR that are more accessible in pSLE B cells were interrogated in the Roadmap/ENCODE database, we found that these regions were enriched for enhancer histone marks H3K4me2, H3K4me3, and H3K27ac (p=0.027) when compared to randomly-generated regions of the genome. This finding is consistent with our previous analysis of SLE genetic risk loci compiled from several GWAS, which are also enriched for the same histone marks (3). Subsequent analysis with ChipEnrich found these DAR to contain consensus binding sites for 42 transcription factors that may be accessible in pSLE patients but not healthy children, including 6 that have been identified in previous GWAS as conferring disease risk: Nr4a1, GFI1b, Eomes, STAT3, Ets1, and Elk1 (37–40). ChipEnrich also identified SUZ12 and EZH2 binding sites to be enriched within the DAR. SUZ12, EZH1 and EZH2 have been shown to regulate recruitment of the transcriptional machinery to transcription start sites of genes important to the activation of the pro-inflammatory response (41). Indeed, using Pscan-ChIP, a web server that finds over-represented transcription factor binding motifs described by motif profiles available in databases such as JASPAR or TRANSFAC, and their correlated sequences in chromatin immunoprecipitation with sequencing experiments (ChIP-seq) (24), several DAR in pSLE B cells displayed enrichment for binding motifs for transcription factors that regulate inflammation, including STAT3, NFkB and PPARg. No difference in accessibility was observed for a control motif, not found to be enriched in either pSLE or healthy B cells. Thus our data identified activation of the pro-inflammatory response in pSLE B cells manifested in changes in chromatin architecture surrounding specific transcription factor binding motifs.

Gene ontology enrichment analysis of 3864 genes expressed within DAR of pSLE B cells, compared against a background of all protein-coding genes (from biomart ensembl), was performed using GORILLA. This revealed 129 different biologic processes present within the DAR, most notably cellular activation in the immune response, regulation of cell proliferation and cellular responses to external stimuli (e.g., interferon).

### 3.2 B cells from pediatric lupus patients show a distinct ATACseq profile when compared with B cells from adult lupus patients

As mentioned above, we downloaded ATACseq data from B cells of adult lupus patients and healthy adults in the GSE118235 dataset, publicly-available on NCBI SRA (14), and applied the pipeline described above for analysis of ATACseq data in our pediatric samples, comparing the results between the two cohorts. Differential peak analysis revealed 12761 DARs, with more regions of chromatin inaccessibility in B cells from adult SLE than pSLE. We also found more peaks located over 100kb from the nearest transcription start site in adult SLE than in pSLE. Peaks located in closer proximity to transcription start sites had less enrichment for transcription factor binding loci in adult lupus B cells than pediatric lupus B cells.

There were 65.2% of the DAR in pSLE patients located within known SLE haplotypes, compared to 39.7% of DAR from B cells of adult SLE patients (p=0.02). Gene ontology enrichment analysis of 1661 genes expressed within adult SLE DARs within known SLE haplotypes revealed 130 different biologic processes, most notably cellular activation in the immune response, and complement activation. The same analysis of overlapping DARs between adult and pediatric SLE within known SLE haplotypes revealed 209 genes belonging to cellular processes important in immune cell activation, including JAK/STAT and TLR signaling, and cellular adhesion (Figures 2e, 2f).

**Figure 2.**
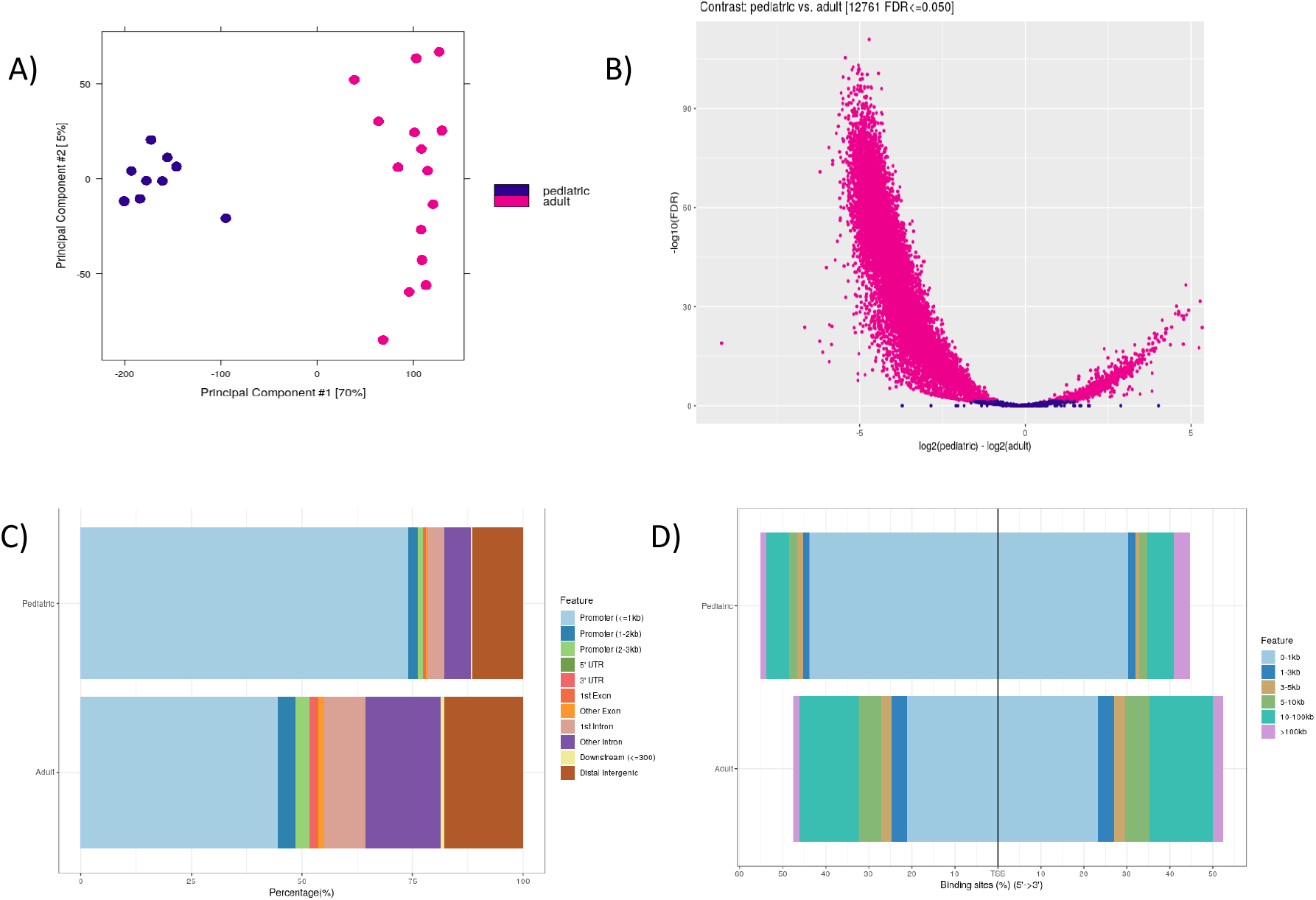
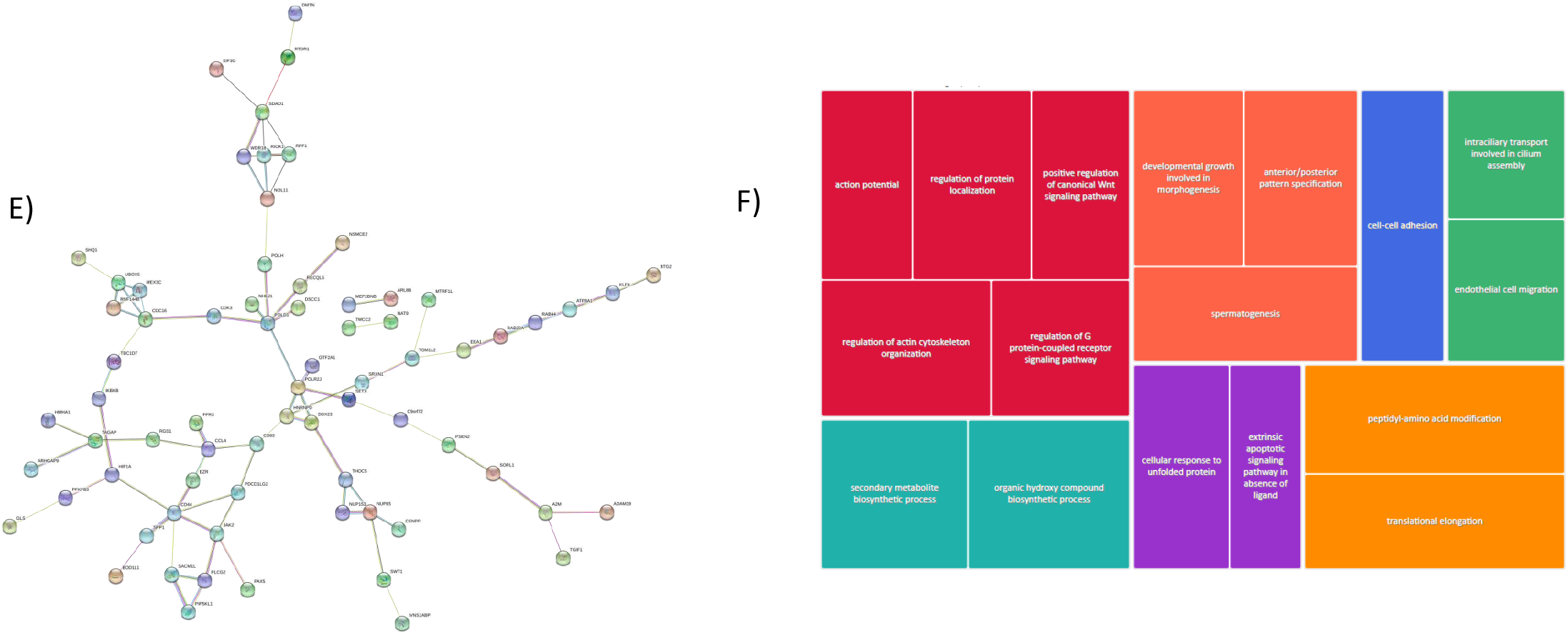
Unique chromatin architecture in pediatric-onset lupus B cells compared to adult lupus B cells. a) Principal component analysis plot showing distribution of pediatric lupus (darker points) and adult lupus (lighter points) samples. b) Differentially accessible regions in pediatric lupus B cells compared to adult lupus B cells (false discovery rate <0.05). Lighter plotted points to the right of 0 on the x-axis indicate increased chromatin accessibility in pediatric lupus patients. Lighter plotted points to the left of 0 on the x-axis indicate potentially less accessible chromatin in pediatric lupus patients. Note that there appears to be more closed chromatin than open chromatin in adult lupus patients compared to pediatric lupus. c) Histogram showing that there are fewer peaks located distally and intergenic in pediatric lupus B cells than adult lupus B cells. d) Histogram showing that peaks found near transcription start sites in pediatric lupus B cells have more enrichment of transcription factor binding loci than adult lupus. e) Protein-protein interaction network of genes interacting with enhancers that overlap differentially accessible binding sites between pediatric and adult lupus B cells (plotted in String). f) REVIGO gene ontology tree map highlighting up-regulated pathways in pediatric lupus but not adult lupus B cells. Shades of grey demonstrate superclusters, and box sizes indicate the strength of the p value (larger size reflects higher statistical significance).

### 3.3 Genes involved in the pro-inflammatory response and cellular adhesion are more accessible in B cells in pediatric lupus than in adult lupus B cells

Using the Integrated Genome Viewer (42), a high-performance visualization tool for interactive exploration of large genomic datasets, many loci with increased accessibility in pSLE patients are located near start sites of their adjacent genes (Figure 3). Fifty-five percent of pSLE DAR located within SLE haplotypes lie within 1kbp of genes involved in cell proliferation, activation of the adaptive immune response, activation of the innate immune response, and protein degradation, e.g., interferon regulatory factor-5 (IRF5) (Figure 3b), STAT4, and TNFAIP3. Polymorphisms in the above genes have been shown to be associated with disease pathogenesis in adult SLE (43,44). STAT4 and IRF5 are important regulators of type I interferons; peripheral B cells in patients with SLE display a prominent, highly reproducible type I interferon gene signature. Similar changes were not observed in the accessibility of IRF5 or STAT4 in B cells from healthy children.

**Figure 3.**
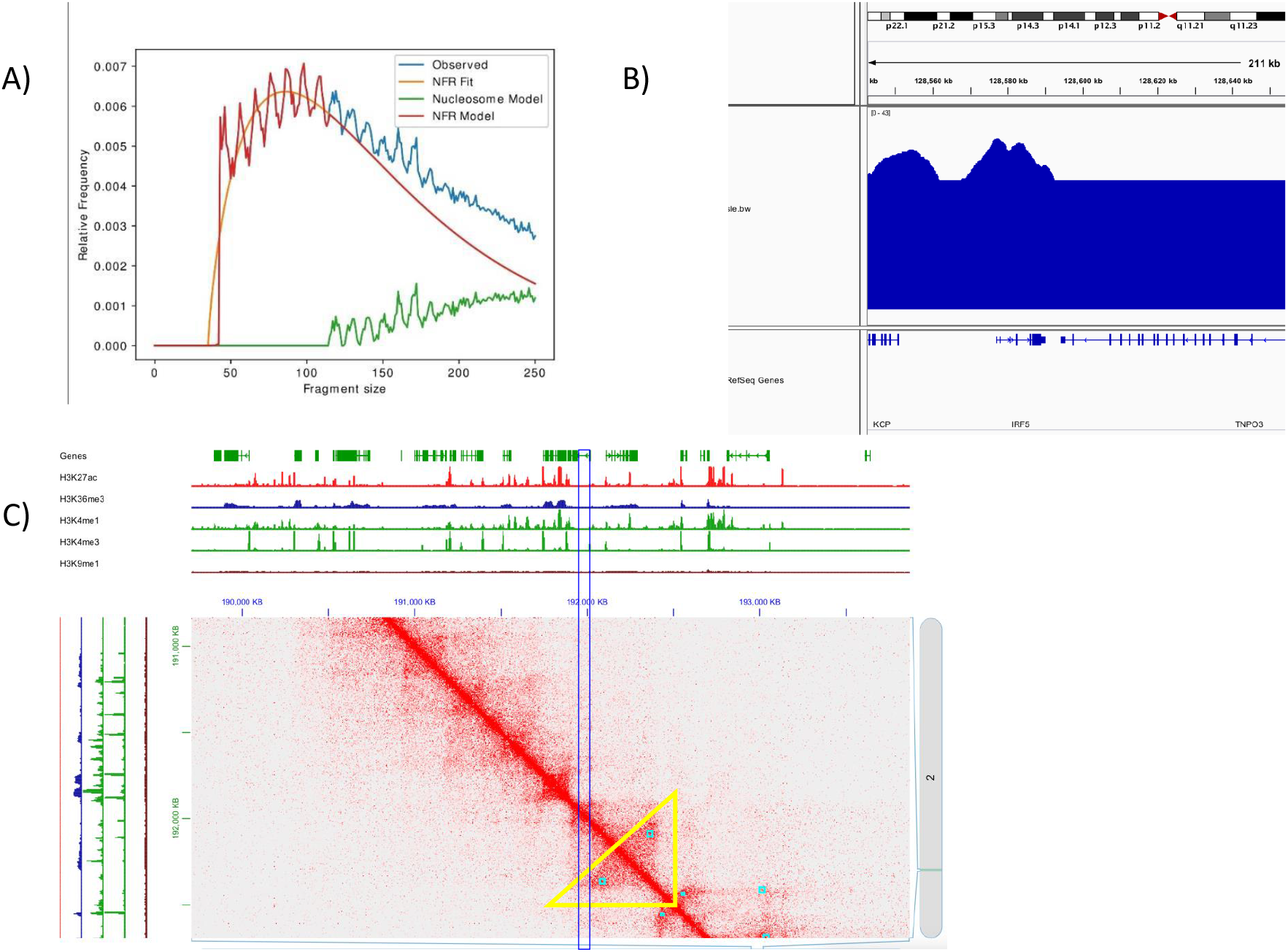
Loci with increased chromatin accessibility in pediatric lupus patients can cluster around transcription start sites of genes important in the pro-inflammatory response, which may then be poised for activation. a) Fragment size distribution as obtained with NucleoATAC. b) Broad peaks from pediatric lupus patients within DAR in SLE haplotype containing IRF5 are clustered near the transcription start site. This haplotype is also enriched for enhancer marks, indicating that the likelihood of transcription of IRF5 is very high in pediatric lupus B cells. c) Representative figure of 3-dimensional chromatin conformation around STAT4. The triangle represents a topographically-associated domain, and the thin rectangle is the location of the SNP used to tag the risk locus. Note that the thin rectangle appears to overlap the topographically-associated domain, indicating close contact with the chromatin loops, and suggesting that STAT4 is poised for activation in pediatric lupus B cells.

Further interrogation of the DAR residing within SLE haplotypes revealed 57 genes, including those important in cell-to-cell adhesion (cadherins) and chemokine-mediated signaling, e.g. CCL4, which have been suggested to play a role in SLE pathogenesis in adults (45,46). Using Juicebox (47), an integrated viewer of Hi-C data, we discovered that several of these genes, including those involved in JAK/STAT signaling, were in close 3-dimensional contact with chromatin loops in B cell lymphoma cell lines, suggesting that they are poised for activation in pSLE B cells (Figure 3c).

## Discussion

We are the first to investigate epigenomic changes in pediatric-onset SLE using ATACseq. In a cohort of 8 children with active, treatment-naïve pSLE patients, who were compared to 5 healthy children, we demonstrated differences in chromatin accessibility in peripheral blood B cells. There were 50.34% of the peaks in pSLE patients located intergenic and distally from promoter sites, indicating that many transcription factors may be acting as distal enhancers to regulate transcription. The DAR from pSLE B cells were enriched for enhancer histone marks. Transcription factor binding motifs for STAT3, NFkB and PPARg were enriched within the DAR. Fifty-five percent of DAR located within SLE haplotypes appear to be within 1kbp of genes involved in cell proliferation, activation of the adaptive immune response, activation of the innate immune response, and cellular adhesion, e.g., CCL4, interferon regulatory factor-5 (IRF5), STAT4, and TNFAIP3.

Our data are similar to ATACseq data in adult SLE B cells: Scharer et al (14) reported increased accessibility in a range of transcription factors involved in B cell activation and B cell differentiation. They also reported differential accessibility between SLE and healthy adult B cells in the STAT4 promoter, leading them to conclude that SLE B cells are epigenetically unique from healthy adult B cells. Our analysis of their raw ATACseq data demonstrated further epigenetic differences between pediatric and adult lupus B cells. Most notably, we found more regions of *inaccessible* chromatin in adult SLE B cells than those from pSLE, suggesting decreased levels of transcription of many associated genes. A larger proportion (65.2%) of the DAR in B cells from pediatric lupus patients were located within SLE haplotypes compared to 39.7% in adult lupus patients, suggesting that more regions of open chromatin in pediatric lupus B cells may be poised for transcription and contribute to disease pathogenesis.

Limitations to our study include a small sample size and the use of total B cells in both pSLE patients and healthy children. We were limited in the starting material for ATACseq, and would not have had enough B cells in each group if we had attempted to perform bulk ATACseq in B cell subsets. However, we acknowledge the importance of studying the B cell subsets individually, as specific subsets have been shown to directly contribute to disease pathogenesis (e.g., plasmablasts). Scharer et al demonstrated feasibility of performing bulk ATACseq in B cell subsets from adult lupus patients (14); however, the amount of starting material to perform bulk ATACseq in each B cell subset would be prohibitive in pSLE patients. Studies are currently underway by our group to determine feasibility of using single-cell ATACseq to evaluate B cell subsets from pSLE patients. Single-cell ATACseq would be the appropriate next methodology to use as it is particularly useful for describing rare, transient populations.

Moreover, we acknowledge that our data would be strengthened by corresponding data from transcriptomes of pSLE B cells compared to B cells from healthy children. Here again, we were limited in our starting material for bulk ATACseq and were unable to perform RNA sequencing in the same samples from treatment-naïve pSLE or healthy B cells. Single cell RNA sequencing has been successfully performed to characterize PBMC from pediatric lupus patients (48), confirming feasibility. Genes important in cellular adhesion and the pro-inflammatory response, including genes contributing to the interferon gene signature, were found to be upregulated in pediatric lupus patients. Several of these genes, including CCL4, IFI44, and ROCK1, were similarly found to reside in DAR described in our study, indicating they are more accessible for transcription in our pediatric lupus cohort. Studies are underway in our laboratory to determine the optimal methodology to investigate genetic variants that cause this rare, pediatric autoimmune disease.

In conclusion, this is the first report describing chromatin architecture in children with pSLE. We demonstrate an epigenetically-distinct profile in pSLE B cells when compared to those from healthy children, healthy adults, and adults with SLE, indicating that pSLE B cells are predisposed for disease onset and development. Differences in chromatin accessibility between B cells in pSLE patients, adults SLE patients, and healthy individuals, particularly in pathways controlling inflammation and activation of the immune response, suggest that transcriptional dysregulation of key players in B cell activation and differentiation plays an important role in the pathogenesis of pSLE.

## Author Contributions

JSHY was involved in study conceptualization and design, data collection, and all aspects of data analysis. JSHY wrote the first draft of the manuscript and performed revisions. KJ and SM performed all wet lab experiments including sample processing, B cell isolation, and building libraries for ATACseq experiments, and helped revise the manuscript. BAE and HW were involved in patient recruitment, sample procurement, and helped draft the manuscript. BD was involved in study conceptualization and design, and helped revise the manuscript. JNJ was involved in study conceptualization, design, and coordination, and wrote the first draft of the manuscript with JSHY. All authors read and approved the final manuscript.

## Acknowledgments

We thank the patients and families for their participation in this study. This work would not have been possible without the bioinformatics expertise of Dr Frank Jenkins (1967-2021). We thank Andrea La Bella and Daniel Lulo for technical expertise and database management. We thank Drs Suhas Ganguli, Mei Ma, Roya Samuels, Minu George, Beth Gottlieb, and Katherine Steigerwald for their clinical care.

## Contributions to the field

Lupus is an autoimmune disease that can affect major organs of the body, and primarily affects girls of child-bearing age. Genetic variations and the environment can alter the human immune response and result in disease. It is well-known that there are differences in gene expression between children with lupus and those without. Epigenomic sequencing techniques used in our study revealed differences in transcription regulation, and increased accessibility, of genes important in activation of the immune system and the pro-inflammatory response in children with lupus, compared to adults with lupus and children without lupus. This provides a more comprehensive picture of what may make children vulnerable to the development of lupus, as well as the differences in response to therapies between patients, and allow us to design new treatments that may be safer and more effective in children than current therapies.

